# Genomic and transcriptomic characteristics of type VI secretion system in *Klebsiella pneumoniae*

**DOI:** 10.1101/2024.01.04.574191

**Authors:** Wanzhen Li, Xiaolan Huang, Dan Li, Xiaofen Liu, Xiaoying Jiang, Xingchen Bian, Xin Li, Jing Zhang

## Abstract

The Type VI secretion system (T6SS) serves as a crucial molecular weapon in interbacterial competition and significantly influences cell-cell interactions. Various bacterial species utilize their T6SSs to execute a multitude of functions, dictated by their ecological niche. However, the characteristics of T6SS in clinical *Klebsiella pneumoniae*, a common opportunistic nosocomial pathogen, have not been fully elucidated. Here, we conducted a genomic analysis of 65 clinical *K. pneumoniae* isolates obtained from patients with varying infections. Genes encoding a T6SS cluster were present in all analyzed strains of *K. pneumoniae*. Strains of identical sequence type (ST) carried structurally and numerically identical T6SS. Our study also highlights the importance of selecting conserved regions in key T6SS genes for effective primer design in PCR identification. We then utilized the predominant ST11 *K. pneumoniae* HS11286 to investigate the effect of knocking out T6SS marker genes *hcp* or *vgrG*. Transcriptome analysis identified a total of 1,298 co-upregulated and 1,752 co-downregulated differentially expressed genes. Additionally, the absence of *hcp* or *vgrG* gene suppressed the expression of other T6SS-related genes within the locus I cluster. Pathway analysis showed that the Δ*hcp* mutant exhibited alterations in transport, establishment of localization, localization and cell processes. Furthermore, interbacterial competition experiments showed that *hcp* and *vgrG* are essential for competitive ability of ST11 *K. pneumoniae* HS11286. This study furthers our understanding of the genomic characteristics of T6SS in *K. pneumoniae* and suggested that the involvement of multiple genes in T6SS of strain HS11286.

**Importance:** Gram-negative bacteria use T6SS to deliver effectors that interact with neighboring cells for niche advantage. *K. pneumoniae* is an opportunistic nosocomial pathogen that often carriers multiple T6SS loci, the function of which has not yet been elucidated. We performed a genomic analysis of 65 clinical *K. pneumoniae* strains isolated from various sources, confirming that all strains contained T6SS. We then used transcriptomics to further study changes in gene expression and effect upon interbacterial competition following knockout of key T6SS genes in ST11 *K. pneumoniae* HS11286. Our findings revealed the distribution and genomic characteristics of T6SS in clinical *K. pneumoniae*. This study also described the overall transcriptional changes in the predominant Chinese ST11 strain HS11286 upon deletion of crucial T6SS genes. Additionally, this work provides a reference for future research on the identification of T6SS in bacteria.

## Introduction

The Type VI secretion system (T6SS) in Gram-negative bacteria is a widely distributed nanomachine that delivers effectors into prokaryotic or eukaryotic cells to attain dominance within a specific niche (1, 2). The T6SS is a phage-like device anchored to the bacterial membrane and typically comprises 13 conserved core components designated TssA-M (3, 4). Hcp (TssD), VgrG (TssI) and PAAR are vital T6SS structural proteins that facilitate the transportation of effector proteins. The latter are transported through the T6SS by either fusing with a T6SS structural component or by non-covalently interacting with one of the core components. In either case, Hcp, VgrG, and PAAR structural proteins are involved. Thus, these three proteins have dual roles, serving as both components and substrates of the T6SS (5–7). Therefore, due to the crucial roles played by these genes in the T6SS, many studies utilize *hcp* and *vgrG* genes as markers to identify the presence or absence of the T6SS cluster (8, 9).

The primary function of T6SS involves interbacterial antagonism encompassing both interspecies and intraspecies competition (10, 11). T6SS effector proteins possess antibacterial activity and may act by lysing crucial macromolecular substances such as bacterial DNA, phospholipids, and peptidoglycans (11–16). However, recent studies have broadened the functional understanding of T6SS effector proteins, including their activities against fungi and eukaryotic cells (17, 18). Bacteria can exploit T6SS to interfere with host cell physiological processes including adhesion, skeletal rearrangement, and evasion of innate immunity to gain additional advantages for colonization, dissemination, and survival (17). Additionally, T6SS is also implicated in metal acquisition (12, 19). Overall, T6SS exhibits diverse functionalities and is integral to microbial adaptive survival.

*Klebsiella pneumoniae* is a significant opportunistic Gram-negative pathogen capable of causing severe nosocomial infections among immunocompromised individuals (20). *K. pneumoniae* typically carries multiple copies of T6SS, with two T6SS clusters being the most common (21). Unfortunately, existing studies examining T6SS in *K. pneumoniae* typically lack clinical strains and, when included, the presence or absence of T6SS determined only by PCR with no genomic information provided. While T6SS systems are known to contribute to bacterial competition, cell invasion and colonization in *K. pneumoniae* (9), major gaps remain in our understanding of how this system contributes to *K. pneumoniae* pathogenesis, necessitating further exploration.

In China the predominant genotype of carbapenem-resistant strains of *K. pneumoniae* (CRKP) is ST11 (78%) (22), and these strains usually contain two T6SS clusters (21). The complete genomic sequences of the clinical ST11 multidrug-resistant strains HS11286 is publicly available and widely used as references in various genetic studies (23, 24). This study investigated the distribution of T6SS in *K. pneumoniae* clinical strains and explored changes in transcription levels upon deficiency of key T6SS genes in HS11286. We first conducted a genomic analysis of T6SS using 65 clinical isolates obtained from patients with varying infections. We then focused on ST11 *K. pneumoniae*, coupling a transcriptome analysis with the deletion of T6SS genes *hcp* and *vgrG*. Our results further our understanding of the characteristics of T6SS in clinical strains of *K. pneumoniae*.

## Materials and Methods

### Strains used in this study

A total of 65 clinical isolates of *K. pneumoniae* Isolates from patients with diverse infections were kindly provided by Institute of Antibiotics, Huashan Hospital. The collection contained isolates collected as part of routine clinical care from 2017 to 2020. ST11 CRKP strain HS11286 (NC_016845) was isolated from a clinical sputum specimen and served as the reference strain for this study (25).

### Whole-genome sequencing and analysis

The entire genome of 65 strain was sequenced using the Illumina NovaSeq 6000 (BIOZERON, Shanghai, China). The raw paired end reads were trimmed and qualitatively controlled by Trimmomatic (version 0.36) (26). *Kleborate* was used to identify the sequence type (ST) and K_locus (http://github.com/katholt/Kleborate) (27). Detailed information of all strains used in the study can be found in Table S1.

### Identification of T6SS in clinical *K. pneumoniae* genomes

All genome sequences were analyzed by BLASTp 2.14.1+ (e-value 1e-5) against T6SS clusters of *K. pneumoniae* in the SecReT6 database, which included HS11286 (NC_016845), NTUH-K2044 (NC_012731), Kp52.145 (NZ_FO834906) and ATCC 43816 (NZ_CP064352) (28). Additionally, the T6SS of KP17-16 (NZ_CP034077.1) identified by SecReT6 was also included in this study. We also designed specific primers of *hcp*, *vgrG*, and *tssM* from HS1286 to perform Polymerase Chain Reaction (PCR) for identifying T6SS cluster in clinical strains. The gene structure of T6SS was visualized using R (29).

### Phylogenetic analysis

A total of 66 isolates was constructed by Snippy v4.6.0 with strains *K. pneumoniae* HS11286 (NC_016845) used as the reference (30). Core SNP alignments were used to remove recombination sites by Gubbins v2.4.1(31). We then used this core genome alignment in Fasttree v2.1.11 to generate maximum likelihood phylogenies (32). Tree visualization was performed using iTOL (33).

### Synteny Analysis

The type i2 T6SS clusters in different ST strains were compared using Easyfig v 2.2.5 with default parameters (e-value 1e-3) (34). The nucleotide sequences and annotations of HS11286, NTUH-K044 and KP17-16, were downloaded from the NCBI RefSeq database.

### Gene deletion and complementation

The gene deletion strains HS11286*-* Δ*hcp* and HS11286*-*Δ*vgrG* were constructed using the λ Red recombinase method (35). For complementation, the DNA sequence of *hcp* and *vgrG* were PCR amplified from HS11286 and cloned into the pHSG398 plasmid (laboratory modification with apramycin resistance). These cloned plasmids were transferred into their corresponding gene deletion mutants by electroporation. All the bacterial strains, plasmids and primers used in this study are listed in TABLES S4–5.

### RNA-Seq and bioinformatic analyses

*K. pneumoniae* HS11286, HS11286*-Δhcp* and HS11286*-ΔvgrG* mutant strains were grown overnight in LB broth (37 °C, 180 rpm). Overnight cultures were diluted 1:100 with fresh LB broth, grown to logarithmic phase (OD_600_ = 0.4–0.6) and centrifuged at 4500 × g for 10 min. RNA extraction and sequencing were performed by Personalbio (Shanghai) using the Illumina Novaseq 6000 platform. Quality assessment and filtration of the raw data were performed by Cutadapt (36). The filtered RNA-Seq reads were mapped to the reference genome (NC_016845) using Bowtie2 (37). Transcript abundance was quantified using the fragments per kilobase per million mapped fragments (FPKM) normalization method. DESeq2 (38) was used for differentially expressed genes (DEGs) analysis using the following conditions: |log_2_FoldChange | > 1, padj < 0.05. Venn diagrams were drawn using jvenn (http://jvenn.toulouse.inra.fr/app/example.html) (39).

Gene Ontology (GO) and Kyoto Encyclopedia of Genes and Genomes (KEGG) functional enrichment analysis were performed to identify which DEGs were significantly enriched in GO terms or metabolic pathways. GO terms and KEGG pathways with a false discovery rate (FDR) of <0.05 were considered significantly altered. All enrichment visualization was performed using R (29).

### Quantitative real-time reverse-transcription polymearase chain reaction analysis

To measure the expression of the T6SS-related genes, total RNA was extracted using the TaKaRa MiniBEST Universal RNA Extraction Kit (Takara, Japan) following the manufacturer’s instruction. Reverse-transcription and qRT-PCR was performed according to the Vazyme’s instructions. The *mdh* gene was used as an endogenous control for all qRT-PCR analyses (40). The relative transcription levels were calculated using the 2^−ΔΔCt^ method.

### Interbacterial competitive assays

Rifampin-resistant *E. coli* EC600 (which lacks a functional T6SS) was used as the prey strain. The predator and prey strains were separately grown overnight in Luria-Bertani (LB) broth at 37° C. Cultures were then diluted 1:100 with fresh LB broth, grown to mid-exponential phase, and resuspended in PBS (OD_600_, ∼0.5; equivalent to ∼10^8^ CFU/mL) after centrifugation at 4500 × g for 5 min. Predator and prey strains were then mixed in a 10:1 ratio, spotted (10 μL) on LB agar containing 100 μg/mL rifampin, and incubated for 5 h or 24 h at 37° C. The mixed spots were harvested, serially diluted in 0.9% saline, and 5 μL of appropriately diluted sample manually plated onto LB agar containing 100 μg/mL rifampin. Plates were incubated for 18–22 h at 37 °C prior to imaging. Additionally, a further 10 μL of appropriately diluted sample was manually plated onto LB agar for viable cell counting. Enumeration was performed manually after 18–22 h of incubation at 37 °C. The prey survival rates were calculated by dividing the CFU (colony-forming units) counts of the output by the CFU of the control.

### Ethics Statement

The Huashan Institutional Review Board approved the sample collection and will waive the informed consent if further studies are conducted using clinical isolates from the culture collection. Personal privacy is not involved in this study.

### Data availability

The genomes sequenced during this study are available under the BioProject accession number PRJNA930978. The RNA sequences have been submitted to the NCBI SRA database under BioProject accession number PRJNA972856.

## Results

### Identification of Type VI Secretion system in clinical *K. pneumoniae* isolates

The distribution of T6SS was evaluated in 65 strains of clinical *K. pneumoniae*. We sequenced clinical *K. pneumoniae* isolates from patients with diverse infections between 2017 and 2020, with HS11286 serving as the reference strain. A phylogeny analysis of 66 clinical *K. pneumoniae* isolates obtained primarily from patients with bloodstream infections (79%; 52/66) was conducted (FIG.1). Of the ST11 and ST15 strains, 88% (15/17) and 88% (15/17) encoded the capsular polysaccharides (CPS) loci KL64 and KL19, respectively. T6SS of clinical *K. pneumoniae* were identification by searching against the existing and predicted T6SS clusters in SecReT6, it was showed that genes encoding a T6SS cluster were present in all analyzed strains. The majority of clinical strains (63/65) were identified to harbor the complete T6SS cluster (Type i2), which encodes conserved T6SS-related genes (FIG.1). However, when PCR was used to identify the presence of *hcp*, *vgrG*, and *tssM* genes, only 32% (21/65) of strains were identified as carrying the T6SS cluster (TABLE S1, supplementary material). The presence of the T6SS gene cluster in all strains precluded identifying any correlation between T6SS and different infection types.

**FIG 1.**
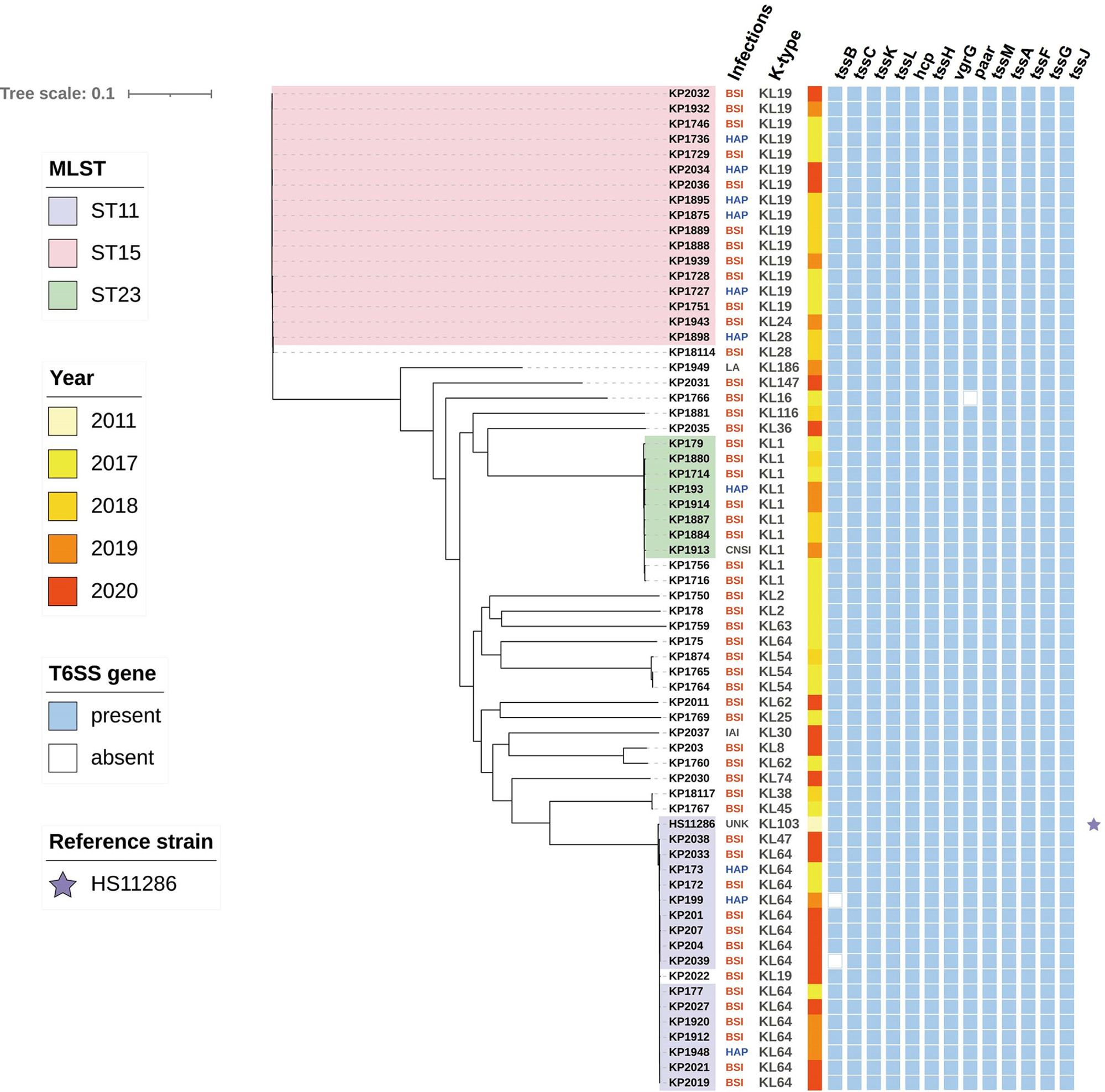
Phylogenetic tree of 66 clinical *K. pneumoniae* strains. The different colors represent different ST types. Column 1 shows the type of infection and column 2 the CPS loci. In columns 3, a colored filled square indicates the collection date. Square colored blue represents the presence of the T6SS-related genes labeled above the figure. BSI: bloodstream infection; IAI: intra-abdominal infection; HAP: hospital acquired pneumonia; CNSI: central nervous system infection; LA: liver abscess; UNK: Unknown.

### General characteristics of T6SS gene clusters in *K. pneumoniae*

Identical sequence type strains were observed to possess identical T6SS clusters (FIG. 2). In this study, all 16 clinical strains of ST11 shared T6SS structures similar to the reference strain HS11286, with each strain harboring two copies of T6SS. All ST23 strains exhibited T6SS structures similar to that of the ST23 strain NTUH-K2044, with two copies of T6SS. And there are minor differences in the gene arrangement compared to HS11286. Additionally, consistent with the ST15 strain KP17-16, each ST15 strains carried three T6SS clusters. Regardless of the number of T6SS copies carried, each strain of *K. pneumoniae* carried a complete T6SS cluster classified as type i2. However, characterizing the T6SS in strains of other sequence types remains challenging due to their limited representation in this study.

**FIG 2.**
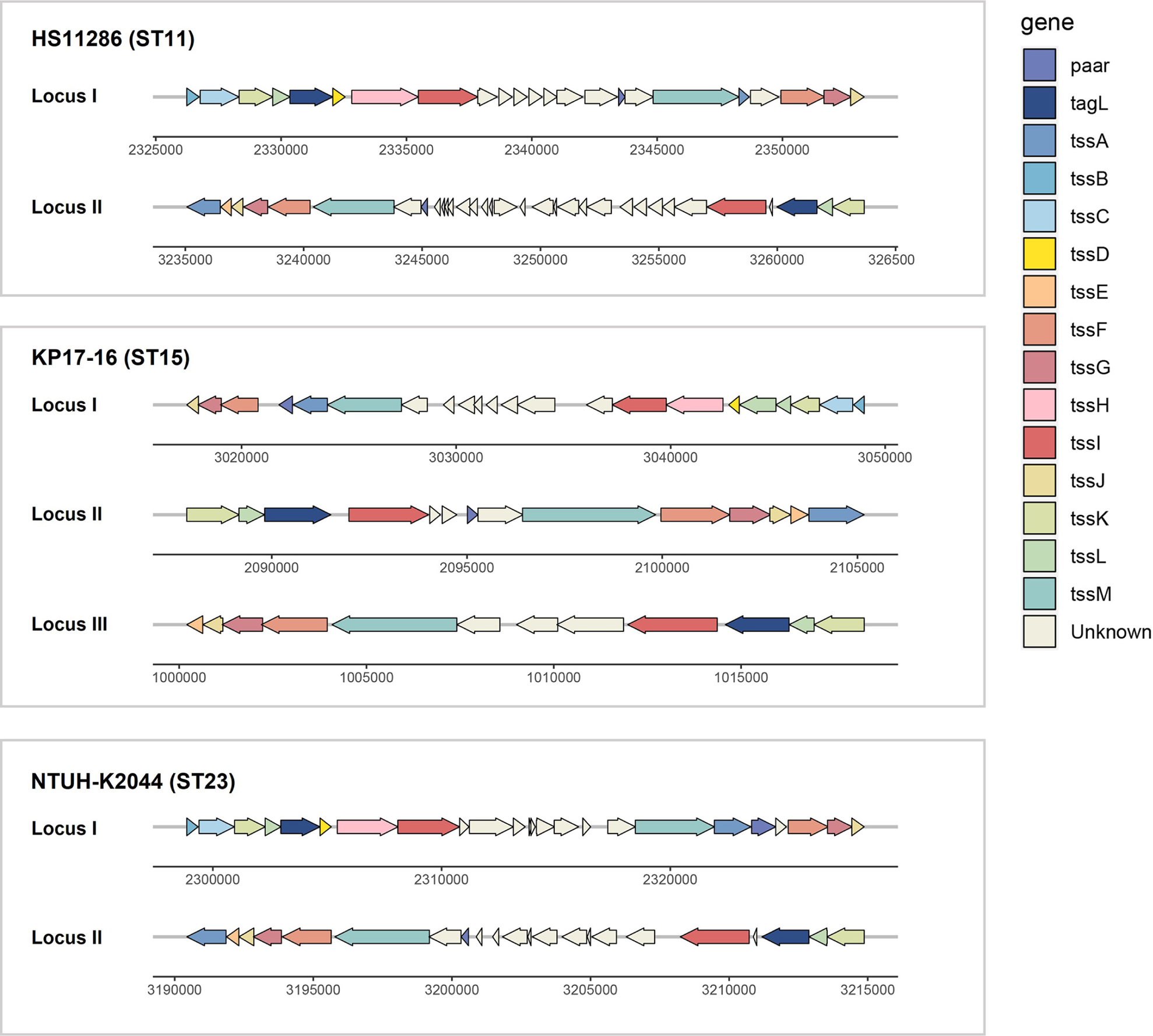
Schematic representation of T6SS clusters in different sequence types of *K. pneumoniae*. Sequence type 11 (ST11) *K. pneumoniae* was represented by strain HS11286, which possesses two copies of T6SS. Sequence type 23 (ST23) *K. pneumoniae* was represented by strain NTUH-K2044, which also possesses two copies of T6SS. Sequence type 15 (ST15) of *K. pneumoniae* contains three copies of T6SS, with strain KP17-16 serving as an example.

### Synteny analysis of type i2 T6SS clusters in *K. pneumoniae*

The complete T6SS cluster in *K. pneumoniae* is classified as type i2 according to the SecReT6 database. We performed a synteny analysis of this type cluster among representative strains of sequence types ST11, ST23, and ST15. Notably, the conserved genes within the T6SS cluster showed over 66% identity (FIG. 3). Comparative analysis of *hcp*, *vgrG*, and *tssM* genes indicated that the hcp gene was highly conserved with nucleotide identities reaching above 99%. Nucleotide multiple alignments (FIG. S2, supplementary material) revealed variability in the *vgrG* and *tssM* genes among different strains. The *vgrG* gene of three strains showed 98% overall identity, but there are significant sequence differences after 1700bp. The *tssM* gene exhibited an overall sequence identity of 97%. However, notable sequence variations are observed specifically within the 1-400 bp region. Strains of the identical sequence type also carried identical *vgrG* and *tssM* genes. therefore, for PCR primer design aimed at detecting *vgrG* and *tssM* genes, selecting a conservative segment of the nucleotide sequence is recommended for optimal specificity.

**FIG 3.**
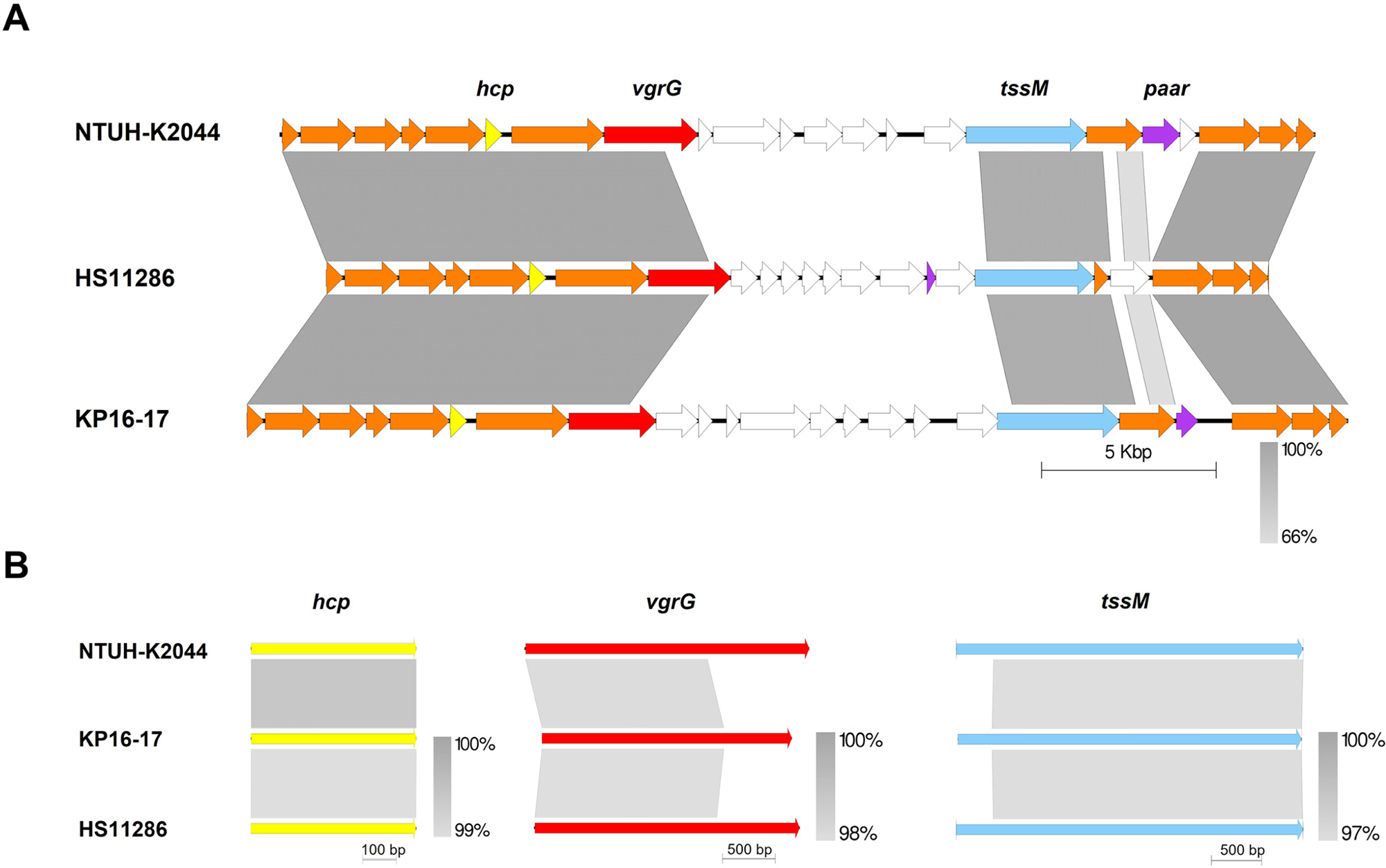
Comparative analysis of the type i2 T6SS cluster in different sequence types of *K. pneumoniae*. (A) The genetic environments of three T6SS obtained from ST23 NTUH-K2044, ST11 HS11286 and ST15 KP17-16 strains are presented as a linear alignment. T6SS-related genes are represented in colored arrows. Bidirectional BLAST hits were illustrated to denote sequence identity ranging from 66%-100%. (B) A comparative gene analysis of *hcp*, *vgrG* and *tssM* among different sequence types of *K. pneumoniae* strains.

### Overview of transcriptomic alterations between mutant strains and WT

To further investigate the role of T6SS in physiological pathways of *K. pneumoniae*, we performed gene knockout of *hcp* or *vgrG* in the ST11 *K. pneumoniae* HS11286 strain and undertook RNA-sequencing (RNA-seq) to compare transcriptomic profiles with the wild-type strain (WT). The number of up- and down-regulated differentially expressed genes (DEGs) is shown in FIG. 4A. RNA-seq revealed a similar number of DEGs in both Δ*hcp* and Δ*vgrG* mutants (3357 *vs*. 3322 total DEGs, respectively). When both mutants were compared to the WT strain, we identified 1298 co-upregulated genes and 1752 co-downregulated genes. The top Gene ontology (GO) analysis with the False discovery rate (FDR) <0.05 were selected for illustration (FIG. 4B). GO analysis revealed only Δ*hcp* mutant DEGs in terms of molecular function (MF) and cellular component (CC).

**FIG 4.**
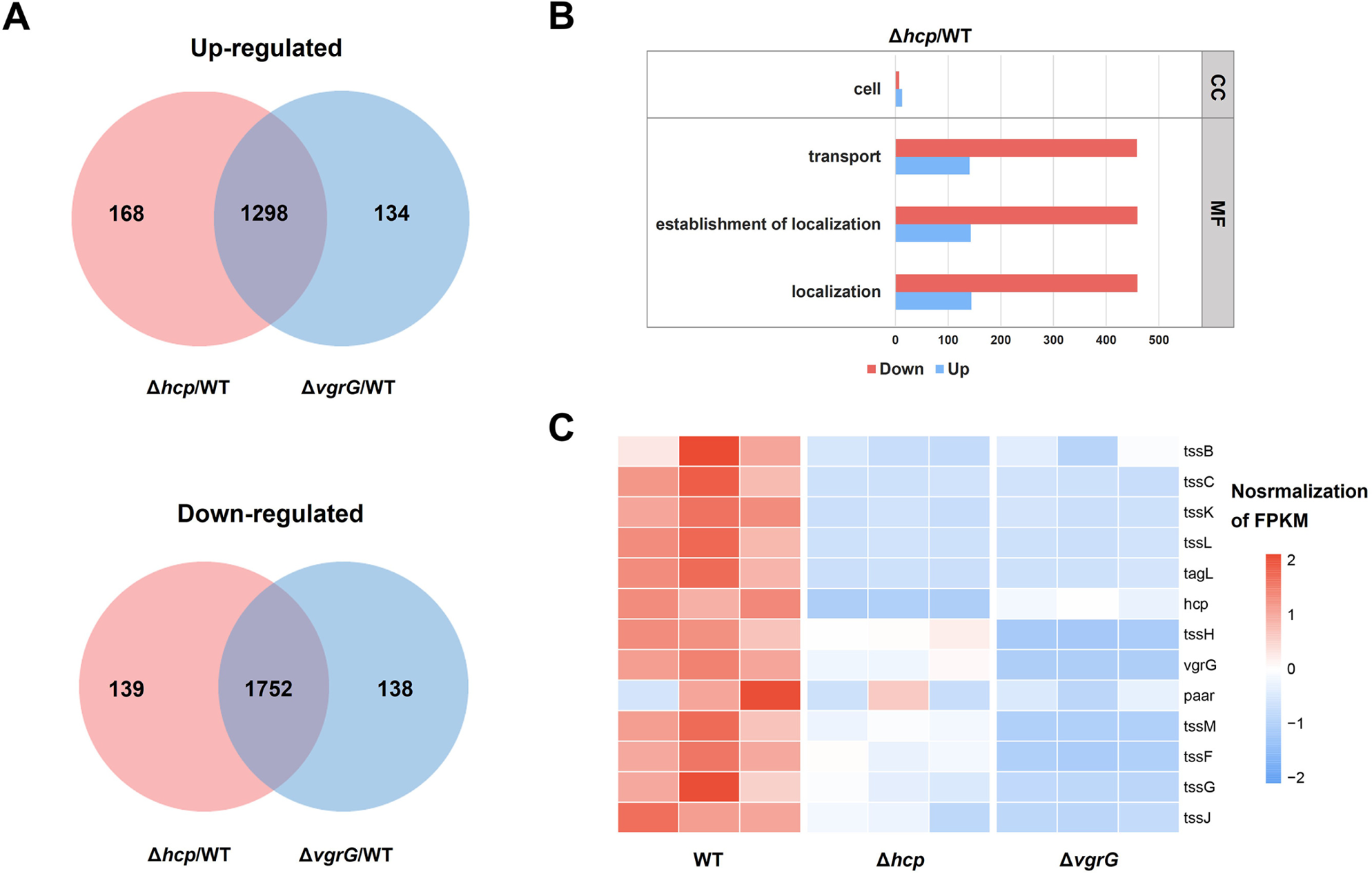
Comparative analyses of the transcriptional response in Δ*hcp* and Δ*vgrG K. pneumoniae* mutants. (A) Venn diagrams showing the number of significantly up-regulated or down-regulated genes (DEGs; |log_2_FoldChange| >1, padj <0.05) in the different *K. pneumoniae* mutants compared to the wild-type (WT). (B) Gene ontology analysis of up- and down-regulated DEGs in Δ*hcp*/WT. CC: cellular component; MF: molecular function. (C) Differential expression of T6SS clusters in Δ*hcp* and Δ*vgrG K. pneumoniae* mutants.

*K. pneumoniae* HS11286 always encodes two copies T6SS clusters. RNA-seq revealed that deletion of either *hcp* or *vgrG* from complete locus I repressed T6SS-related genes within locus I (FIG. 4C). In locus I, only *tssC*, *tssK*, *tssL,* and *tagL* were down-regulated with knockout of *hcp*, while the remaining genes exhibited no significant changes (TABLE S2). In the Δ*vgrG* mutant, *tssC*, *tssK*, *tssL*, *tagL*, *tssH*, *tssM*, *tssF*, *tssG*, and *tssJ* were down-regulated. RT-qPCR analysis showed expression of *hcp, vgrG* and *tssM* of both mutants are consistent with the above transcriptome results (FIG. S3).

### Hcp and VgrG are required for interbacterial competition in *K. pneumoniae*

The Hcp and VgrG proteins are key structural elements of the T6SS. To investigate their contributions to the functioning of *K. pneumoniae*, we assessed the antibacterial activity of corresponding mutants against *Escherichia coli* EC600in a T6SS-dependent manner. A significant (100-fold) reduction in growth of EC600 was observed when co-cultured with the HS11286 WT compared to the Δ*hcp* and Δ*vgrG* mutant strains (FIG. 5). When the mutants were complemented with a plasmid containing either the *hcp* or *vgrG* gene, antibacterial activity against EC600 was restored. Colony count analysis indicated that following 5-h co-culture with EC600, both Δ*hcp* and Δ*vgrG* mutants exhibited diminished *in vitro* competitive ability compared to the WT, although the difference did not reach statistical significance (FIG. S4). However, after 24 h of co-culture, the survival rate of EC600 was significantly higher in the presence of the mutants compared to the WT (WT vs. Δ*hcp P*=0.0034, WT vs. Δ*vgrG P*=0.0066).

**FIG 5.**
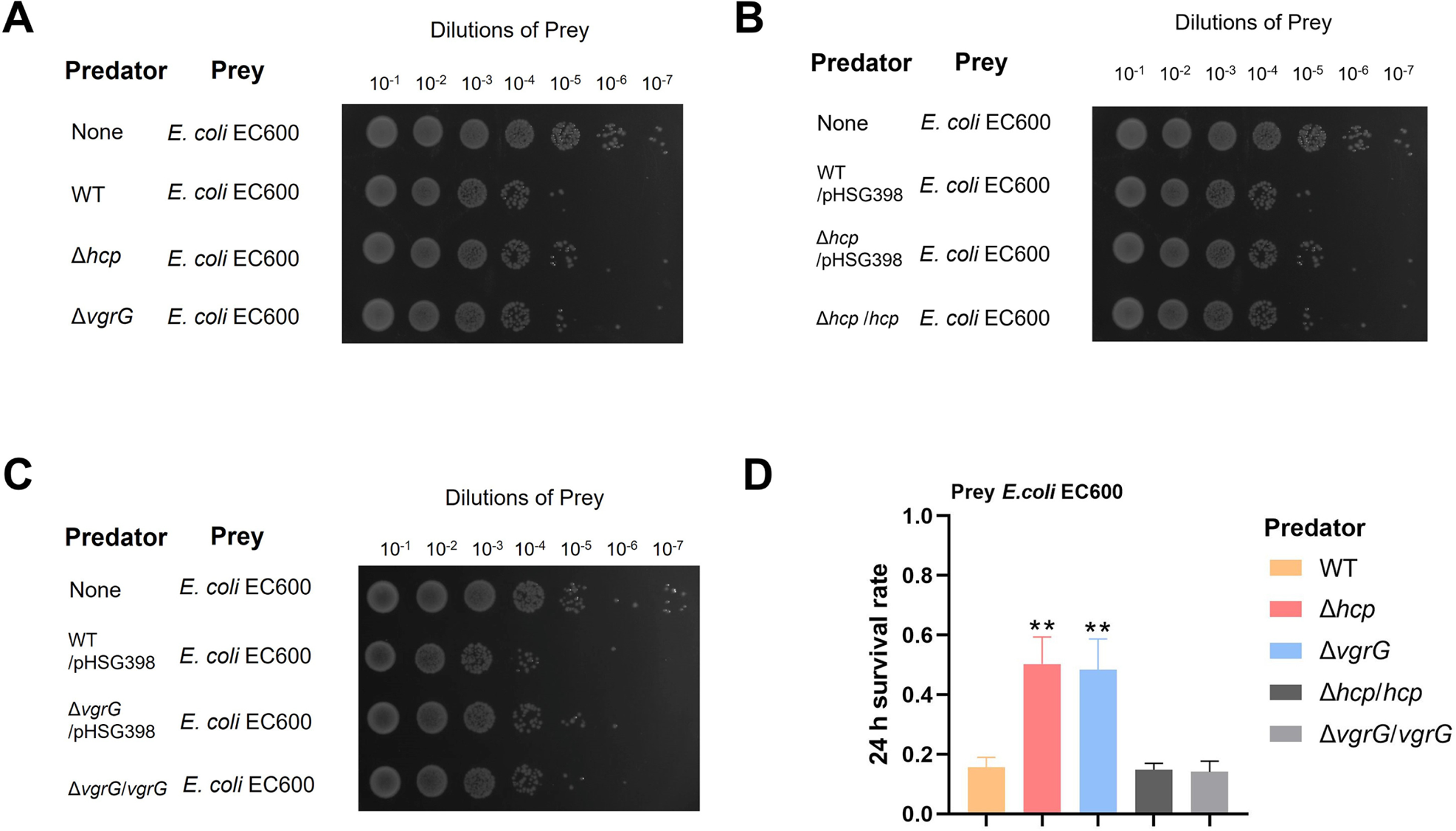
Interspecies killing by *K. pneumoniae* HS11286 in a contact-dependent manner. (A–C) Interbacterial competitive growth assays. Surviving *E. coli* EC600 after 24 h of coincubation with: (A) HS11286 (WT) and the Δ*hcp* and Δ*vgrG* mutants; (B) The WT containing the empty vector (WT/pHSG398), the Δ*hcp* mutant containing the empty vector (Δ*hcp*/pHSG398), and the Δ*hcp* mutant complemented with pHSG398-*hcp* (Δ*hcp*/*hcp*); (C) WT/pHSG398, the Δ*vgrG* mutant harboring pHSG398 (ΔvgrG/pHSG398), and the Δ*vgrG* mutant harboring pHSG398-*hcp* (Δ*vgrG*/*vgrG*). A control lacking any *K. pneumoniae* strain (None) was also included. Recovered mixtures were plated onto Luria-Bertani agar supplemented with 100 μg/mL rifampicin. (D) Survival rates of prey strain EC600 following 24 h of coincubation with the associated predator strain (*K. pneumoniae* HS11286 (WT), Δ*hcp* mutant, Δ*vgrG* mutant, *hcp* and *vgrG* complemented strains). The data represent the means of 3 independent trials. \*\**P* <0.01 by two-tailed Student’s *t*-test (Δ*hcp* mutant or Δ*vgrG* mutant compared

## Discussion

The T6SS plays a pivotal role in bacterial remodeling due to its intra- and inter-species antibacterial activity (41–44). As the predominant genotype of *K. pneumoniae* varies depending on the region of the body in which it is found and the type of infection (9), we studied the distribution of T6SS in *K. pneumoniae* strains isolated from bloodstream, pneumonia, and other types of infections. The genome sequencing results revealed the presence of T6SS clusters in all 65 clinical *K. pneumoniae* isolates. Furthermore, strains of identical sequence type (ST) generally exhibit a similar construction of T6SS cluster. The predominant ST11 *K. pneumoniae* in China always contain two copies of T6SS. Similarly, strains of ST23 also carry two T6SS clusters, whereas strains of ST15 harbor three. Given the considerable size of the T6SS cluster, which spans approximately 20-30 kilobases, the identification of certain T6SS genes through draft genome sequencing may be susceptible to bias. Since only 32% of the strains in this study were identified by PCR as carrying T6SS clusters, and PCR failed to ascertain the copy number of T6SS within a strain. Consequently, for accurate identification of T6SS in strains, priority should be given to complete genome sequencing, followed by the draft genome. When employing PCR, it is imperative to select the conserved region of the pivotal T6SS gene for primer design.

Bacteria adapt their T6SS apparatus to secrete effector proteins with specific functions depending on the environmental niche (41, 45). *K. pneumoniae*, notably the prevalent domestic ST11, typically carries two T6SS clusters. The two key T6SS structural proteins, Hcp and VgrG, serve dual roles as both components and substrates of T6SS and are released extracellularly upon T6SS activation (46, 47). Furthermore, Hcp and VgrG also function as “carriers” for the secretion of effector proteins (6). In this study, we found that deletion of either *hcp* or *vgrG* genes in T6SS locus I led to a range of similar DEGs alterations in *K. pneumoniae* HS11286. But in the Gene Ontology (GO) analysis, only the deletion of *hcp* exhibited notable changes in transport, establishment of localization and localization of the Molecular Function (MF) category, as well as the changes in cell of the Cellular Component (CC) category. Transcriptomic analysis also showed that deletion of *hcp* or *vgrG* genes inhibited expression of certain genes in the complete T6SS gene cluster. Given that the secretion process of T6SS is cyclical whereby Hcp and VgrG proteins are involved in assembly of a sheath that will contract (‘fire’), perforating the membrane of target cells and delivering effector proteins before ultimately being disassembled prior to the next cycle (46), it remains to be clarified whether deletion of these proteins affects both the assembly and secretion processes.

T6SS is widely regarded as a contact-dependent secretory system. Our results showed that knockout of *hcp* or *vgrG* hampers the ability of *K. pneumoniae* to compete when co-incubated with another bacterial species. Given T6SS is a contact-dependent bactericidal mechanism, these results also suggest that prolonged inter-strain contact (24 h in our experiments) allows *K. pneumoniae* to exert its T6SS competitive advantage. However, this does not mean that T6SS is directly involved in the pathogenesis of *K. pneumoniae*. Instead, it is possible that T6SS can eliminate potential microbial competitors to benefit its own colonization. For example, *Yersinia pseudotuberculosis* uses T6SS-3 to secret nuclease effector protein Tce1 to kill other bacteria and facilitate gut colonization in mice (48).

In conclusion, the presence of T6SS was confirmed in all 65 clinical isolates of *K. pneumoniae* examined. Strains of identical sequence type (ST) carried structurally and numerically identical T6SS. This study also provides a foundation for future identification of T6SS clusters in bacteria. In the absence of sequencing results, designing primers based on the conserved nucleotide sequences of key genes can provide a more accurate determination of the presence or absence of T6SS. Within the carbapenem-resistant *K. pneumoniae* strain HS11286, *hcp* and *vgrG* were confirmed as essential proteins for T6SS activity. The deletion of both genes induced similar DEGs changes, and inhibited expression of the locus I T6SS gene cluster. This study further enhances our understanding of the characteristics of T6SS in *K. pneumoniae.* Future studies examining the function of T6SS in *K. pneumoniae* will further enhance our understanding of its adaptive capabilities in response to the environment.

## Supplemental Material

**TABLE S1** Background information of 65 clinical *K. pneumoniae* strains used in this study.

**TABLE S2** Prevalence of T6SS-related genes in *K. pneumoniae* strains.

**TABLE S3** T6SS-related genes expression was changed in mutants compared to the wild-type (WT) strain.

**TABLE S4** Strains and plasmids used in this study.

**TABLE S5** Primers used in this study.

**FIG S1** Multiple sequence alignment among *tssM* and *vgrG* genes

**FIG S2** RT-qPCR T6SS-related gene expression levels in Δ*hcp* and Δ*vgrG* mutants compared with the wild-type (WT) strain.

**FIG S3** Interspecies killing by *K. pneumoniae* HS11286 in a contact-dependent manner.

## Acknowledgments

We gratefully acknowledge Jian Li and Phillip Bergen for their assistance and helpful discussions. This study was supported by the National Natural Science Foundation of China (82173896) and Shanghai Municipal Health Commission (20234Y0012).

